# Designing, optimizing, and assessing modular functional near-infrared brain imaging probes using an automated software workflow

**DOI:** 10.1101/2021.04.06.438705

**Authors:** Morris Vanegas, Miguel Mireles, Qianqian Fang

**Affiliations:** Northeastern University, Department of Bioengineering, 360 Huntington Avenue, Boston, MA, USA, 02115

**Keywords:** functional near-infrared spectroscopy (fNIRS), head gears, modular probe, brain sensitivity, probe optimization, analysis

## Abstract

**Significance:** The exponential growth of research utilizing functional near-infrared spectroscopy (fNIRS) systems has led to the emergence of modular fNIRS systems composed of repeating optical source/detector modules. Compared to conventional fNIRS systems, modular fNIRS systems are more compact and flexible, making wearable and long-time monitoring possible. However, the large number of design parameters makes designing a modular probe a daunting task.

**Aim:** We aim to create a systematic software platform to facilitate the design, characterization, and comparison of modular fNIRS probes.

**Approach:** Our algorithm automatically tessellates any region-of-interest using user-specified module design parameters and outputs performance metrics such as spatial channel distributions, average brain sensitivity, and sampling rate estimates of the resulting probe. Automated algorithms for spatial coverage, orientation, and routing of repeated modules are also developed.

**Results:** We developed a software platform to help explore a wide range of modular probe features and quantify their performances. We compare full-head probes using three different module shapes and highlight the trade-offs resulting from various module settings. Additionally, we show that one can apply this workflow to improve existing modular probes without needing to re-design or re-manufacture them.

**Conclusion:** Our flexible modular probe design platform shows promise in optimizing existing modular probes and investigating future modular designs.

## 1 Introduction

Functional near-infrared spectroscopy (fNIRS) is an emerging neuroimaging technique to non-invasively measure brain activity using non-ionizing light.^1^ Similar to functional magnetic resonance imaging (fMRI),^2^ fNIRS utilizes near-infrared (NIR) light to detect brain activation by measuring the associated hemodynamics. Its use of minimal equipment has positioned fNIRS as an imaging modality of interest to address the challenges of conventional neuroimaging techniques, including non-invasive continuous monitoring, limited spatiotemporal resolution, and the immobility of the user during experiments.^3^ It has shown great promises for safe and long-term monitoring of brain activity and is increasingly used in studies such as behavioral^4^ and cognitive neurodevelopment,^5–8^ language,^9,10^ psychiatric conditions,^11,12^ stroke recovery,^13^ and brain-computer interfaces.^14–16^

Despite an exponential growth in the number of applications^17,18^ and publications^3^ in recent years, many fNIRS systems still employ fiber-based, cart-sized instrumentation^19^ that place limits on both channel density and the use of fNIRS in natural environments, hindering our progress towards understanding the brain. Although fiber-based high-density^20^ and portable^21^ fNIRS systems have been demonstrated, the use of fragile fiber optics cables, stationary external source/detector units,^22, 23^ and the need for individual and specialized headgear for specific tasks have motivated the fNIRS community to investigate more flexible modular and fiber-less designs.^24,25^

The modular fNIRS architecture is based on utilizing identically fabricated elementary optical source and detector circuit (modules) used as repeating building blocks to form a re-configurable probe.^24^ This modular architecture offers significantly improved portability, scalability, flexibility in coverage, and lowered fabrication cost.^24^ By avoiding the use of fragile optical fibers, modular fNIRS systems permit the use of light guides to directly couple light sources and detectors to the scalp, significantly reducing the signal loss due to fiber coupling. The lightweight and compact modules also make wearable fNIRS and continuous monitoring in mobile environments possible.^3, 26^ In addition, the ability to use both intra-module (within a single module) and inter-module (source and detector on different modules) channels allows for high density probes with varying source-to-detector separations (SDS) that increase measurement density and tissue depth sampling, resulting in enhanced signal quality, and easy removal of physiological noise.^27^

Despite these benefits, the task to design a modular fNIRS probe can quickly grow in complexity as the number of modules involved increases. Even for a single module, determining and optimizing a large array of parameters, including module shapes, optode numbers, and optode locations can become a daunting task.

Aside from these core parameters, published studies have also placed emphases on mechanical, ergonomic, safety, usability, optoelectronic, and communication considerations.^24^ For example, mechanical features such as optical coupling and encapsulation must be considered alongside ergonomic considerations such as comfort, weight, and robustness. Additionally, the use of high density light sources in such modular probes bring about additional safety considerations, such as heat dissipation, driving voltage, and battery life. Morever, optoelectronic considerations arise from the use of specialized optodes with narrow emission bandwidths, high gains, low noise, and fNIRS-optimized wavelengths. Not only are these specialized optodes more expensive due to their niche applications and characteristics, they also require more complex control electronics for driving optodes and acquiring data. With such dense coverage, complex encoding strategies such as frequency^28^ multiplexing become a necessity to obtain high quality data acquisition to achieve sufficient spatial and temporal resolution. Finally, while previously reported modular fNIRS systems often employ daisy-chain communication protocols to connect multiple modules on a single bus,^29–33^ the design of physical inter-module connections,^34^ the synchronization method between modules,^24^ and the transfer of acquired data become increasingly complex.

To address some of these factors, a number of fNIRS data analysis packages exist.^35–37^ However, most of these tools focus heavily on motion artifact correction,^35^ quality assessment,^37^ and statistical analysis of channel data.^35,36^ While some tools exist that assist in probe design,^38–41^ they usually assume user-provided optode positions and do not have the capability to tessellate, connect, or re-orient the locations of modules. There is a lack of a systematic approach to analyze, optimize, and compare modular fNIRS systems since the final probe is dependent on many design- and experimental-specific factors.

Here, we are not proposing a solution to address every design challenge mentioned above, rather, we report a framework that provides a simplified and systematic approach to describe and automate the characterization of modular fNIRS probes. This allows researchers to analyze and improve existing modular probes, design new fNIRS modules, as well as create region-specific fNIRS headgears using optical modules. The entire workflow has been implemented into an open-source, MATLAB-based toolbox called Modular Optode Configuration Analyzer (MOCA^42^) to provide the fNIRS community with a tool to guide design decisions as they develop future modular systems. MOCA supports a list of commonly used module shapes, user-defined optode layout and region-of-interest (ROI) coverage, and can produce quantitative performance metrics such as distributions of source-detector (SD) separations, sensitivity maps, and spatial multiplexing groupings. These performance metrics also allow comparisons between different designs of modular probes. Limited manual editing capability is also provided to allow designers to delete, rotate, and tessellate modules for added flexibility.

The remainder of the paper is outlined below. In Section 2, we discuss the relevant design considerations when developing a modular probe using MOCA. We specifically focus on the parameterization of the modules and headgears formed based on these modules, processes required to assemble modules into functional probes, and related performance metrics for system characterization and comparisons. In Section 3, we demonstrate MOCA’s capability in designing full-head probes using a variety of module shapes and compare their trade-offs regarding channel density, SD separations, and average brain sensitivities. Furthermore, we utilize MOCA to showcase potential improvements to published fNIRS probes by optimizing module orientations, spacing, and staggering layouts. In the Section 4, we discuss MOCA’s assumptions and limitations.

## 2 Methods

A diagram showing the overall design process of a modular fNIRS system is shown in Fig. 1. Specifically, the three parts describing MOCA’s workflow are 1) the design parameters describing a single module design, 2) the processes used to assemble the modules into a probe, and 3) the derived performance metrics used to characterize the resulting probe. MOCA starts with the definition of essential module parameters (such as geometry and optode layout) and probe-related constraints (such as ROI and maximum SD separation), shown in the left column in Fig. 1, applies those parameters in an automated probe-generation process (center column in Fig. 1), and derives quantitative characteristics of the resulting probe (shown in the right column in Fig. 1). Arrows in Fig. 1 define dependencies between the derived performance metrics and the input parameters. For example, in order to calculate the probe’s channel distribution, one must define the module geometry, ROI, and optode layout design parameters.

**Fig 1.**
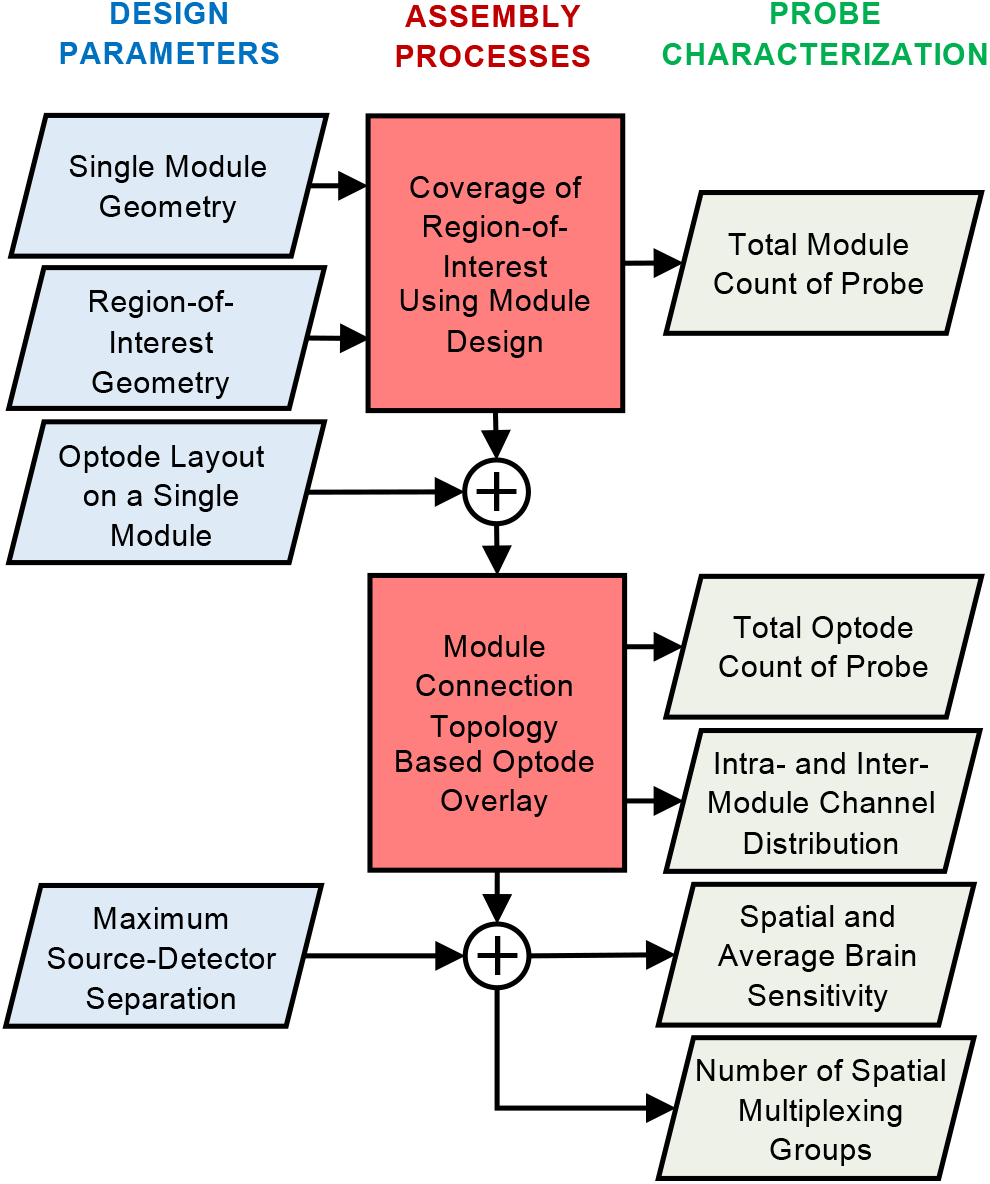
Workflow of design parameters (left column; blue) used in processes (center column; red) to produce specific probe characteristics (right column; green). Probe characteristics are organized top to bottom from least complex (two parameters needed) to most complex (four parameters needed). Arrows trace how parameters are used to derive specific probe characteristics.

### 2.1 Essential design parameters of fNIRS modular probes

The basic building block of a modular probe is an fNIRS module. It is typically in the form of an optoeletronic circuit made of a rigid^29,30, 33,43^ or rigid-flex^44,45^ substrate with on-board light sources, optical sensors, auxiliary sensors, microcontrollers and other communication electronics. A modular probe is subsequently constructed by replicating and interconnecting multiple identically modules, therefore, the design decisions regarding the module parameters are highly important and directly impact the functionalities and restrictions of the resulting probe.

#### 2.1.1 Single module geometry

The shape of a module is one of the key parameters when designing a modular system. In published literature, simple polyhedral shapes, especially equilateral polygons (square, hexagon, etc), are typically used due to their simplicity to fabricate, analyze, and tessellate a target ROI. It is also possible to design probes that combine multiple polygonal shapes, such as a combination of hexagonal and pentagonal modules. Such hybrid-shape modular systems may bring advantages in tessellating curved surfaces, but they also requires more complex analysis. MOCA supports a number of built-in module shapes including three equilateral polygons (triangle, square, hexagon). In such cases, one is only required to define the module edge size as the only shape parameter. One should be aware that a small-sized module requires a large number of boards to cover a given area, thus, resulting in higher fabrication cost and higher complexity in assembly and analysis. Moreover, a small module size also limits the maximum intra-module SDS. Shorter SD separations are known to be more sensitive to superficial tissues rather than brain activities. On the other hand, a small-module size provides better probe-to-scalp coupling when a rigid-board based module is used. MOCA provides limited support for user-specified arbitrary polygonal modules, defined by a sequence of two-dimensional (2-D) coordinates. However, subsequent analyses of these user-defined modules only use the bounding box of these polygons for simplified tessellation and interconnecting.

#### 2.1.2 Target regions-of-interest

An ROI refers to the area of the scalp directly above the cortex for which brain activities are expected to occur.^46^ For simplicity, here we focus on designing probes based on the coverage of a 2-D ROI. For generality, in MOCA, an ROI geometry is specified as a closed polygon made of a sequence of 2-D coordinates. Users need to specify at least three Cartesian coordinates to define a closed ROI. In the future, MOCA can potentially be expanded to support using three-dimensional (3-D) surfaces as ROIs through the use of 3-D surface tessellation tools, such as Iso2Mesh^47^ mesh generator and 3-D photon transport modeling tools such as NIRFAST^48^ and Monte Carlo eXtreme^49^ (MCX).

#### 2.1.3 Optode layout within a single module

Optode layout refers to the spatial arrangement of optical sources and light sensors within the boundaries of a single polygonal module. In MOCA, each source and detector position is defined by a set of discrete 2-D coordinates relative to the module’s center. The 2-D coordinates define the center of the active area of the light-emitting-diode (LED), laser, or photo detector. For simplicity, the physical dimensions of the optodes as well as the size and location of electronic components needed to drive each optode are not considered. The SD separations between all combination of SD pairs are automatically calculated based on the optode positions.

#### 2.1.4 Maximum source-detector separation and maximum short separation channel

MOCA also considers the maximum SD separation (*SDS_max_*) as a key design parameter. Typically, *SDS_max_* is determined by the signal-to-noise ratio (SNR) of the detected signal.^50^ A large SDS has low detector sensitivity due to the exponential decay of the light intensity as SDS increases. This maximum separation limits the number inter-module channels that emerge from a particular tessellation of modules over an ROI. By default, MOCA considers any SDS below 10 mm to be a short-separation (SS) channel. This threshold can be manually changed to fit any specific optode performance or probe application. MOCA uses 30 mm as the default *SDS_max_*.^51,52^ MOCA bounds the SD range by the SS channel threshold and the *SDS_max_*.

### 2.2 Probe assembly processes

A modular probe is constructed when multiple modules are arranged to form a non-overlapping coverage of the ROI area. The final probe is dependent on the tessellation (the number of modules and the spacing between them) and the orientation of each individual module in the probe.

#### 2.2.1 Automated module spatial tessellation

MOCA provides an automated process to tessellate any module shape over a user-defined ROI, which is generally known as the “tiling” problem in computational geometry.^53^ Here, a “complete tessellation” refers to the tiling of an ROI using a single module shape without overlapping or leaving a gap in coverage. Each of the three built-in polygons (triangle, square, hexagon) have the ability to completely tessellate a 2-D area.^54^ MOCA performs the tessellation by first tiling the module shape along a horizontal axis starting at the lowest vertical coordinate of the ROI until the width of the row composed of adjacent modules is wider than the width of the corresponding segment of ROI the row is tiled over. It then repeats this row-generation process until the height of all the rows combined is larger than the maximum height of the defined ROI. This dimension comparison in both axes accounts for module shapes with non-vertical and non-horizontal sides. For irregular module shapes, MOCA uses the maximum width and maximum height of the defined polygon as the a bounding box to create a tiling grid of the module over the ROI. Using the maximum width and height of the ROI as a guide for tiling ensures the full ROI is covered. Although MOCA automatically offsets and flips the three equilateral polygon shapes to prevent gaps, irregular module shapes have inherent gaps between modules when tessellated. Additionally, MOCA accepts manually defined tessellations by reading a sequence of coordinates defining the center of modules to specify each individual module’s location within the ROI. Following tessellation, each module is assigned a unique index and an adjacency matrix is constructed to represent which modules are next to one another.

Limited manual editing capability is provided in MOCA to allow users to modify individual modules and extend the flexibility of probe creation. Users can change probe spacing, the minimum distance between adjacent modules in all directions. Additionally, a module can be manually deleted from the tessellation to allow the probe to more closely follow the boundaries of the ROI or better represent intentional empty spaces in the probe. When individual modules are removed from the probe, the adjacency matrix is automatically re-calculated from the resulting probe.

#### 2.2.2 Module orientation and routing

Module orientation refers to the rotation of the module along the normal direction of the ROI plane. In a “complete tessellation” of the three equilateral polygon shapes, MOCA appropriately flips and translate modules to prevent gaps and overlaps. For tessellations of irregular shapes, each module is simply placed in the same orientation as it was originally defined. After automatic probe generation, MOCA allows the user to manually change the orientation of individual modules based on their assigned indices. For asymmetric optode layouts, changing the module orientation alters the SDS of inter-module channels, resulting in different performance metrics. MOCA alerts the user if a manual re-orientation results in overlapping modules in the probe.

Additionally, MOCA creates a single sequential path to connect all modules to form a linear data communication bus, referred to as the “routing” process. In such a path, all modules are connected and every module is visited exactly once—a classic problem known as the Hamilton path^55^ in graph theory. In most configurations, a Hamilton path is not unique and computing such path is known to be an NP-hard problem, i.e. problems that do not have a polynomial complexity when the node number grows. However, due to the limited module numbers commonly used in an fNIRS probe, for simplicity, an exhaustive search of the adjacency matrix can typically identify all Hamilton paths in a given tessellation with no more than a few minutes of computation. For any computed path, MOCA then orients each module based on the angle of a vector defined by the center of the oriented module and the center of the following module in the path. The orientation angle is relative to the horizontal axis.

### 2.3 Performance metrics

#### 2.3.1 Total module and optode counts

Based on the module design and tessellation, MOCA computes the total number of modules, *n_m_*, needed to cover the ROI. In addition, MOCA also outputs the total number of sources (*n_s_*) and detectors (*n_d_*) of the final probe. Module and optode counts are performance metrics used as markers for the estimated cost and usability of a probe. All modules, sources, and detectors of an assembled probe are given unique identifiable index numbers (*m_i_*, *s_i_*, and *d_i_*, respectively).

#### 2.3.2 Channel distribution

For any assembled probe, MOCA generates histograms of the SD separations for all combinations of SD pairs. Particularly, it outputs separately the distribution of inter- and intra-module channels that are below the *SDS_max_* previously defined by the user. These channel distributions aid the user to optimize the probe design based on the targeted application and population. For example shorter channels are more applicable to infant populations. Additionally, MOCA outputs channel density, a metric commonly used for fNIRS probe bench marking. Channel density is defined as the number of channels, *n_channels_*, divided by the area of the ROI.^24^ Furthermore, MOCA can provide a spatial plot overlaying channels on the assembled probe, allowing for visual inspection of low channel density areas within the probe.

#### 2.3.3 Average brain sensitivity

Brain sensitivity (*S_brain_*) refers to the magnitude of the measurement signal change at a detector given a unitary perturbation of optical properties of brain tissue.^56^ A higher *S_brain_* value suggests the probe is more sensitive to the anticipated brain activation. It is calculated from the spatial probability distribution of photons scattering through complex tissue as they travel from the source to the detector.^57^ Although 3-D head and brain anatomies are widely used to compute *S_brain_* in 3-D, the high computational cost and added complexity of considering 3-D brain anatomy limits the software’s usability and efficiency. For simplicity, MOCA uses a five-layer slab model consisting of tissue imitating the scalp, skull, cerebral spinal fluid (CSF), white matter (WM), and gray matter (GM) to determine the spatial sensitivity profile for each SD pair in a probe.^58^ The thickness of each tissue layer in the slab is set to the average thickness of that tissue type computed using the top half of a tetrahedral brain model.^59^ We define the brain region as the combination of gray matter and white matter tissues. The optical properties and resulting thicknesses for each tissue type are summarized in Table 1.

**Table 1.**
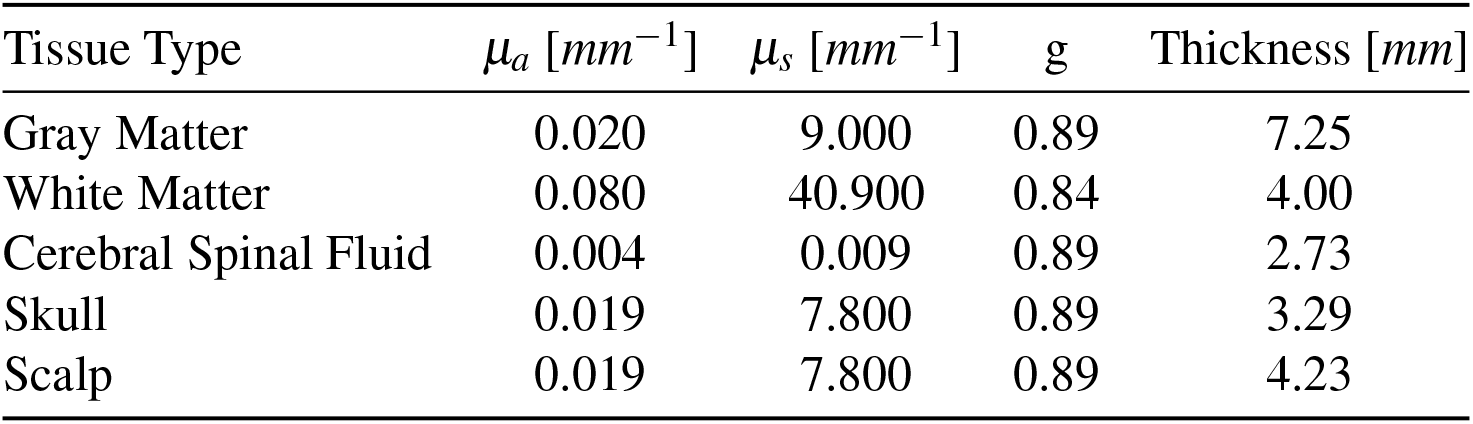
Optical properties used in the slab model for calculating brain sensitivity based on Fang *et al.*^60^ The thickness of each layer is derived by dividing the total tissue volume by the tissue’s surface area from a tetrahedral five tissue brain model.^59^ The absorption coefficient, *μ_a_*, is the average path a photon will travel in the medium before being absorbed. Similarly, the scattering coefficient, *μ_s_*, defines the average path length of photons before a scattering event. Anisotropy, g, is a unit less measure of the amount of forward direction retained after a single scattering event.

For each SD pair in the assembled probe, 3 × 10^8^ photons are simulated using our in-house 3-D Monte Carlo photon transport simulator, MCX,^49^ using a pencil beam source and a single 1.5 mm radius detector placed at the surface of the slab at its corresponding SDS. In a voxelated grid, *S_brain_* is defined as a ratio dividing the region-wise summation of the sensitivity matrix in each brain tissue region by the summation of the entire sensitivity matrix for each source–detector separation,^57^ i.e.

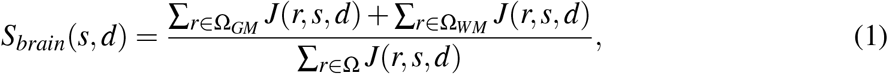

where the sensitivity matrix, also known as the Jacobian (*J*), is computed using the adjoint Monte Carlo method.^61^ In addition to *S_brain_*, MOCA also calculates the average brain sensitivity for the entire probe, 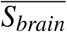, based on all the SD separations above the SS threshold. SS channels are excluded in the calculation of 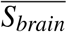 because, by definition, they are designed to only sample superficial layers.^57^

#### 2.3.4 Spatial multiplexing groups

The density of assembled modular probes may impact the probe’s temporal sampling rate when illuminating each source sequentially. Spatial multiplexing is an encoding strategy that can potential accelerate the data acquisition by simultaneously turning on multiple light sources at the same time. Because of the high attenuation of light in the head and brain tissues at large separations, for a given detector, MOCA can ignore the cross-talk of light sources that have separations greater than *SDS_max_* and assign sources into spatial multiplexing groups, or SMG, so that all sources within an SMG can be turned on simultaneously. Notably, unlike frequency multiplexing, spatial multiplexing does not require extra energy-intensive hardware or post-measurement separation of combined signals.

The search for the SMG starts by randomly specifying a source position as the seed; a circle of radius *SDS_max_* centered at the seed position is drawn and a random source outside of this circle that is at least 2 × *SDS_max_* away is picked, and the above process repeats until no additional source can be found. Once an SMG is identified, a new source that does not belong to any existing SMG is selected as the new seed for the next SMG and the above process repeats until every source is allocated. The total number of spatial multiplexing groups, *n_SMG_*, depends on the tessellation of the module over the ROI as well as the choice of the seed position. As with channels, the *n_SMG_* are for a single wavelength, thus, when estimating the total sampling rate of the probe using dual-wavelength sources, the control unit must cycle through each group twice (once for each wavelength).

In addition to *n_SMG_*, MOCA calculates the spatial multiplexing ratio (SMR), defined as *SMR* = *n_s_/n_SMG_*. This ratio is interpreted as the acceleration factor of the data acquisition speed when using spatial multiplexing. For example, for a 20-source probe, an *n_SMG_* of 5 can accelerate the data acquisition by a factor of *SMR* = 20/5 = 4 fold.

## 3 Results

In this section, we first validate the *S_brain_* derived from a simplified five-layer slab model against previously published atlas-based *S_brain_* results.^56^ Then we demonstrate how MOCA can be used to characterize and compare full-head probes composed of different choices of elementary module designs. Lastly, we show examples of using MOCA as an investigational tool to improve existing designs by altering tessellation settings such as module orientation and the distance between modules.

### 3.1 Slab-based brain sensitivity corresponds with atlas-based sensitivity

Fig. 2 shows *S_brain_* calculated using our five-layer slab model at SD separations ranging from 1 to 60 mm in 1 mm increments (blue line). We also overlay full-head average *S_brain_* and standard deviation at 20, 25, 30, 35, and 40 mm separations from a previously published study^56^ using the Colin27 atlas.

**Fig 2.**
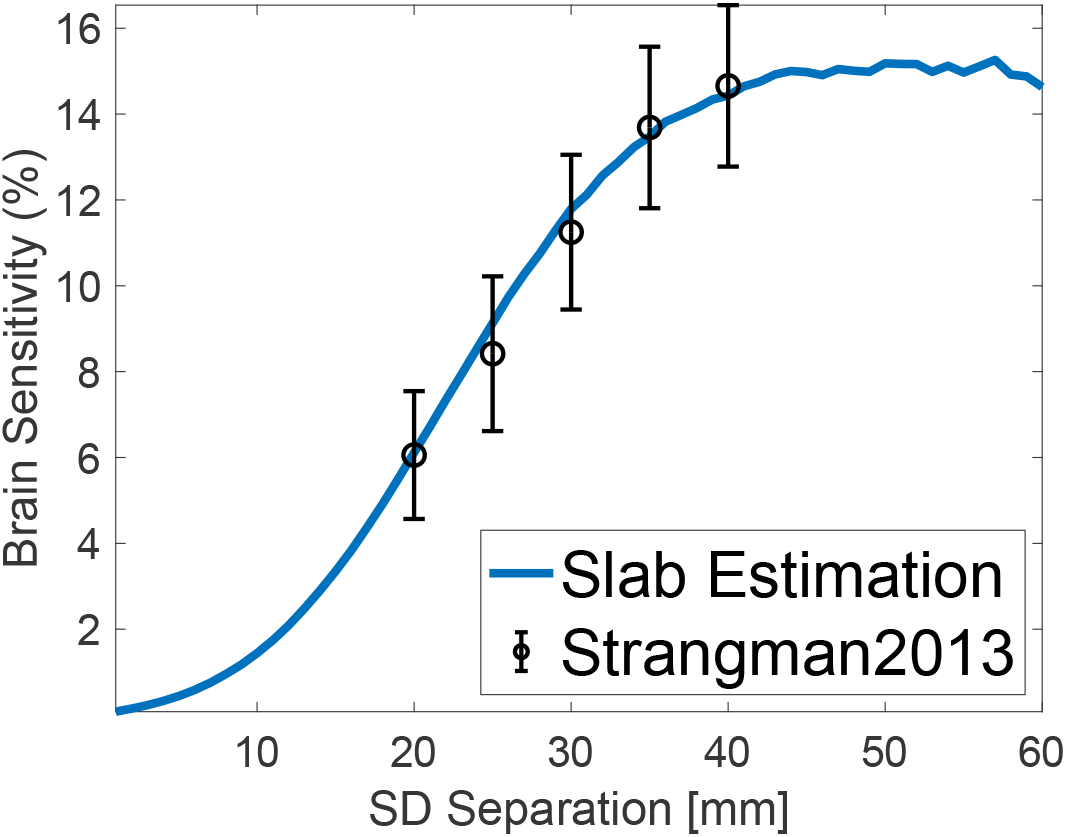
Results comparing brain sensitivity derived from finite slab models and atlas-based models.^56^ The blue line shows calculated brain sensitivity based on a five-layer slab model for SD separations from 0 to 60 mm in 1 mm increments. Overlaid in black are the brain sensitivity results calculated from an atlas across nineteen locations in the international 10-20 system.^56^

Simulations on a five-layer slab model show an increase in *S_brain_* as SDS increases. Additionally, *S_brain_* for SD separations below 10 mm is less than 1.17%. At 20, 25, 30, 35, and 40 mm separations, the maximum difference between the atlas-based and slab-based *S_brain_* values is less than 0.6%. Fig. 2 demonstrates that using 2-D approximation of the ROI and a layered brain structure provides a reasonable trade-off between accuracy and computational efficiency, especially for high density probe characterization.

### 3.2 Comparison between sample modules of various shapes

MOCA provides quantitative metrics that allow the comparison of a wide range of fNIRS module designs to investigate the effects of design parameters on probe characteristics. As a showcase, here we report the results from a comparison of three equilateral module shapes (square, hexagon, and triangle) with the same optode layout tessellated over a 200 × 200 mm ROI, derived from the average surface area of the top half of an adult male head.^62^ Square^29–31^ and hexagonal^32,33,43^ fNIRS modules have been extensively studied in literature and are chosen here for a quantitative comparison. While an equilateral triangle has not been reported in published module designs, we include it here because of the potential suitability for better tessellation of a 3-D surface in future extensions. With this comparison, we want to demonstrate both scalabilities of MOCA in analyzing full-head probes and how probe characteristics of modules with the same optode layouts vary as the module shape changes.

As mentioned above, MOCA automatically tessellates the target ROI using the module geometry and assigns each module an index number. If not considering within-module optode locations, for both square and hexagon modules, only translation is needed to completely cover a region. For the triangle shape, MOCA rotates every other triangle and its optodes 180 degrees to fill the ROI without leaving any gaps. No other orientation changes are made for this comparison. Probe spacing is set to zero. The default SS threshold is set to 10 mm and the *SDS_max_* is set to 30 mm. To avoid simultaneously changing multiple parameters and only focus on module shape, an identical optode layout made of two sources and two detectors is used in all three module designs in this example. The edge-length of the square is set to 33.33 mm, determined by the average length of three previously reported square-shaped module designs.^29–31^ The edge-length of the hexagon and triangle is set to 20.68 and 50.65 mm, respectively, calculated to achieve the same area as the squared module. The three module designs as well as the tessellation of the hexagon-based probe over the ROI are shown in Fig. 3. The derived characteristics for each of the three sample probes are summarized in Table 2.

**Table 2.** Summary of quantitative metrics automatically derived by MOCA when tessellating the three elementary module shapes over a 200×200 mm region of interest.

| Row | Probe Characteristics | Square-based probe | Hexagon-based probe | Triangle-based probe |
| --- | --- | --- | --- | --- |
| 1 | Total modules [N] | 36 | 42 | 40 |
| 2 | Total optodes [N] | 144 | 168 | 160 |
| 3 | Total channels [N] | 324 | 405 | 496 |
| 4 | Intra-module channels [N] | 180 | 237 | 336 |
| 5 | Inter-module channels [N] | 144 | 168 | 160 |
| 6 | % of channels that are inter-module [%] | 55.56 | 58.52 | 67.74 |
| 7 | Average brain sensitivity [%] | $7.52 \pm 1.95$ | $6.50 \pm 2.44$ | $8.83 \pm 3.10$ |
| 8 | Average intra-module brain sensitivity [%] | $6.44 \pm 2.10$ | $6.44 \pm 2.10$ | $6.44 \pm 2.10$ |
| 9 | Average inter-module brain sensitivity [%] | $8.82 \pm 0.00$ | $6.54 \pm 2.66$ | $9.94 \pm 2.86$ |
| 10 | Spatial multiplexing groups [N] | 9 | 8 | 13 |

**Fig 3.**
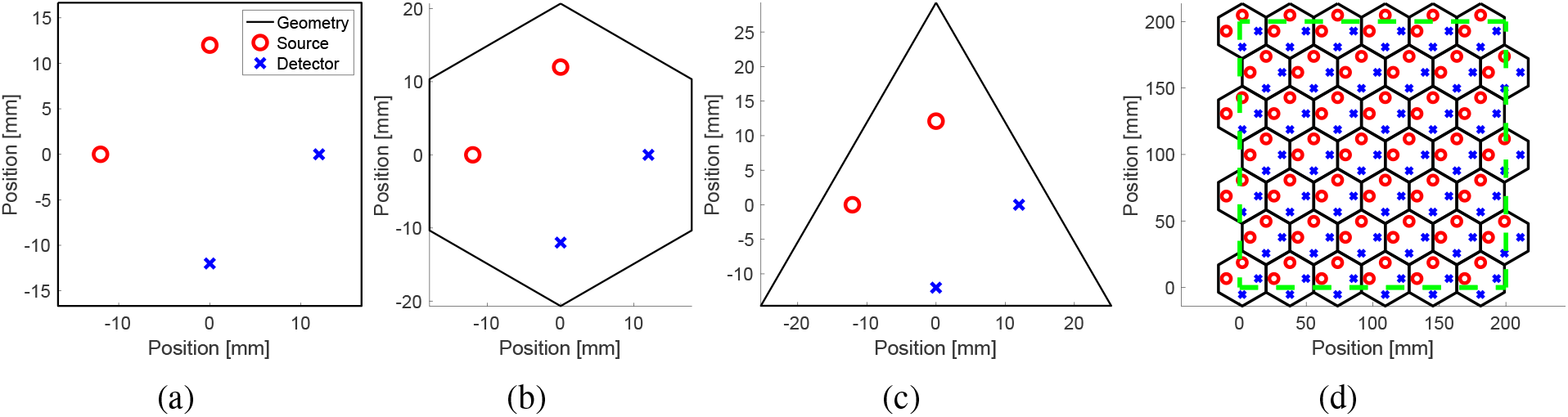
Elementary module designs used in a full-head comparison. (a), (b), and (c) show the perimeter of the square, hexagon, and triangle-based module designs, respectively. The optode layout of all three shapes is identical. Red circles represent sources while blue crosses represent detectors. (d) Tessellation of the hexagon module over an ROI. The dashed green line outlines the 200×200 mm ROI.

#### 3.2.1 Effect of module shape on channel separation distributions

Fig. 4 shows a histogram of the SD separations of the full-head (200×200 mm area) probe composed from the three selected module shapes. Table 2 shows that the number of modules needed to completely cover the ROI varies for each shape due to MOCA’s tessellation algorithm that ensures full coverage of the ROI area using shapes with non-vertical and non-horizontal edges (Fig. 3d). Since each module utilizes the same optode layout, the intra-module channel distributions (blue bars in Figs. 4a, 4b, and 4c) are simply scaled by the total numbers of modules needed to completely cover the ROI. The SDS of inter-module channels are dependent on the module shape, resulting in varying inter-module channel distributions between all three probes (orange bars in Figs. 4a, 4b, and 4c).

**Fig 4.**
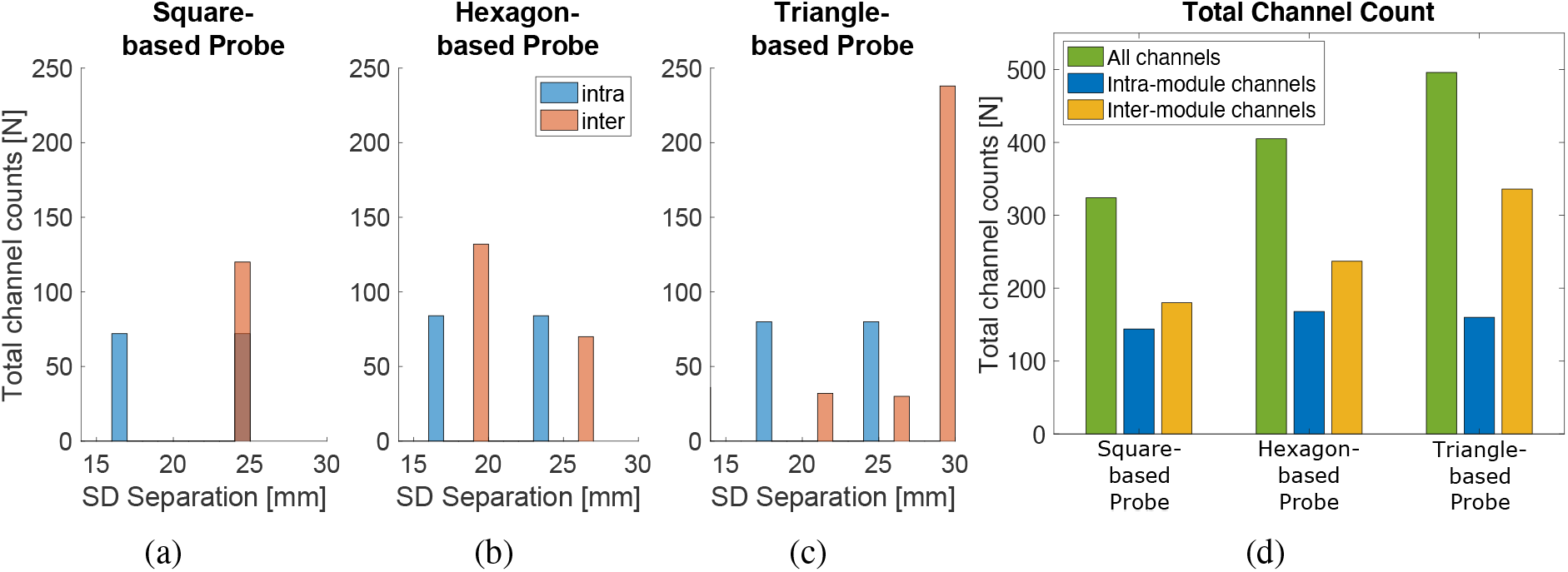
Channel distributions and total channel counts resulting from the tessellation of the three elementary module shapes over a 200×200 mm region of interest. (a-c) Resulting intra- and inter-module channel distributions for square, hexagon, and triangle module-based probes. (d) The total channel count of each probe grouped by intra- and inter-module channels.

For this particular example, the triangle-based probe reports both the highest number of total channels (Fig. 4d) and the largest SD separations of all three tessellated probes (Fig. 4c). The hexagon-based probe appears to have the shortest inter-module channels (Fig. 4b). Due to its symmetry and given the *SDS_max_* setting, the square-based probe happens to have all SD separations at 24 mm. Notably, the triangle-based probe adds the most inter-module channels, almost twice the number of intra-module channels (Fig. 4d), while also requiring two fewer modules than the hexagon-based probe (Table 2, Rows 1-5). Fig. 4d also shows that the number of inter-module channels is greater than the number of intra-module channels for all three probes.

#### 3.2.2 Combining intra- and inter-module channels for brain sensitivity

The 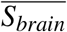 values derived from the three probe designs, grouped by intra-module channels, inter-module channels, and or all channels, are summarized in Fig. 5. Only channels above the SS threshold and below the *SDS_max_* are used. Despite having the fewest total channels (Table 2, Row 3), the square-based probe results in a higher 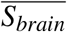 than the hexagon-based probe. For the square- and triangle-based probes, the use of inter-module channels increases the probe’s 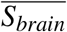 as compared to simply using intra-module channels alone. For the hexagon-based probe, 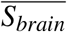 computed using only intra-module channels is similar to that when using only inter-module channels (6.44% vs 6.54%). Due to having the same optode layout, the intra-module 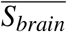 is the same for all three probes.

**Fig 5.**
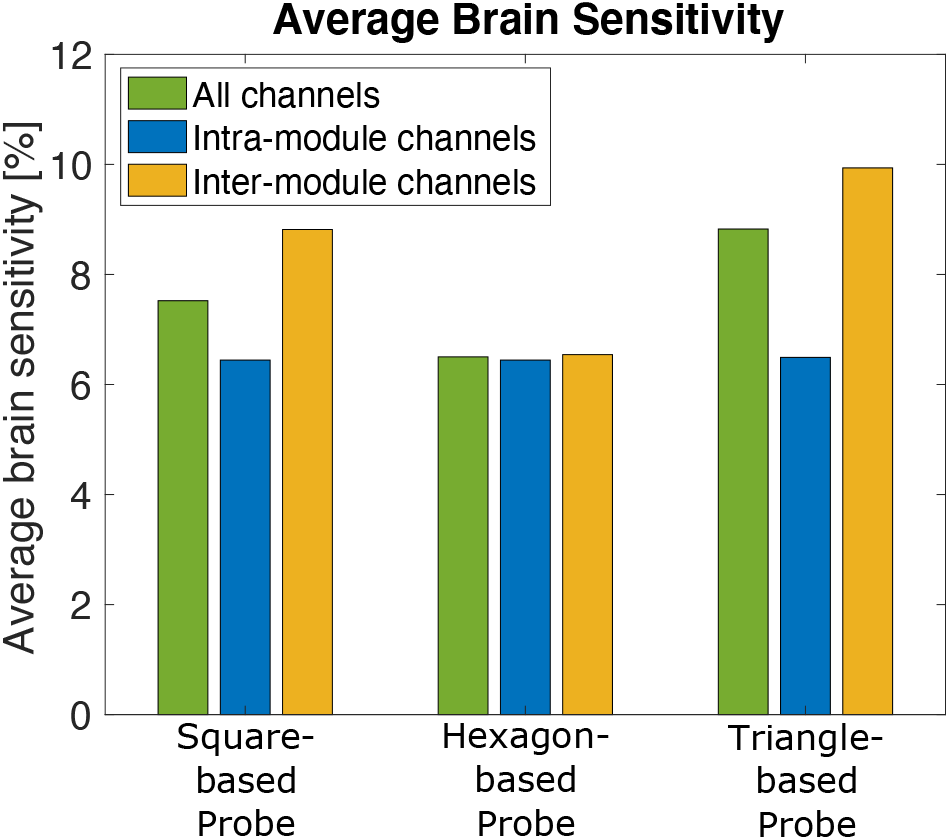
Resulting average brain sensitivity organized by intra- and inter-module channels for square-, hexagon-, and triangle-based probes tessellated over a 200×200 mm region. Short-separation channels are excluded in all calculations.

#### 3.2.3 Effect of module shapes on the number of spatial multiplexing groups

The total *n_s_* compared to the *n_SMG_* arising from the tessellation of each module over the ROI are compared in Fig. 6a. Fig. 6b overlays the first SMG over the triangle-based full-head probe. The total number of sources for the square-, hexagon- and triangle-based probes are 72, 84, and 80, respectively. Using the *n_SMG_* for each probe (Table 2, Row 10), the SMR (the ratio between *n_s_* and *n_SMG_*) is 8, 10.5, and 6.15 for the square-, hexagon-, and triangle-based probe, respectively. This result indicates that the hexagon-based probe’s sampling rate can benefit the most when using group-based spatial multiplexing.

**Fig 6.**
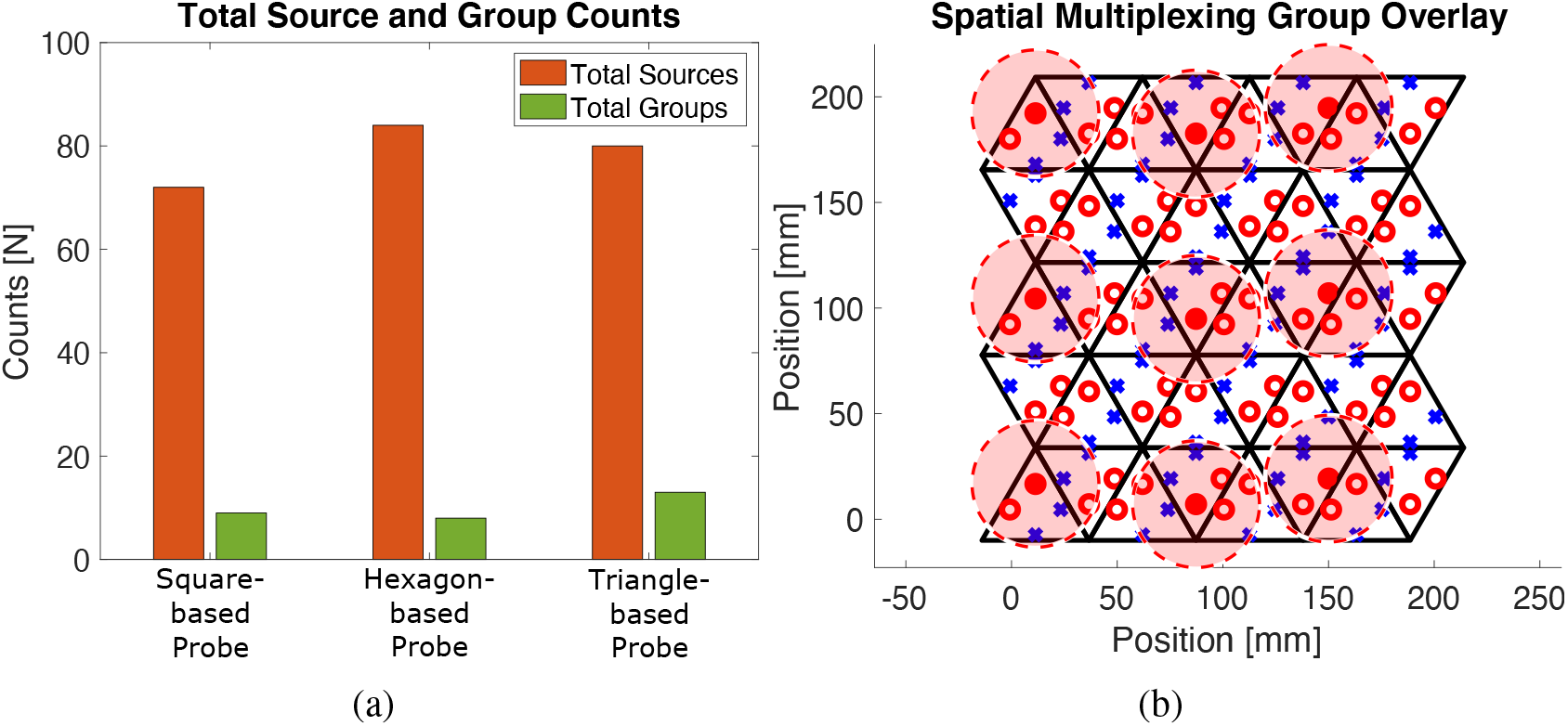
Spatial multiplexing group results from the tessellation of the square-, hexagon-, and triangle-based probes. (a) Comparison of total number of sources (orange) and total number of spatial multiplexing groups (green). (b) The triangle-based module tessellation with sources (red circles) and detectors (blue crosses). The dashed red circles indicate the “effective” region (30 mm radius) of each of the nine sources in the first spatial multiplexing group. The nine sources turned on simultaneously in this group are indicated by filled in red circles.

### 3.3 Improving existing probes through probe layout alterations

The ability to compute probe characteristics from basic design parameters allows users to explore probe tessellation alterations and improve existing probes using MOCA. Here, we alter published examples to demonstrate how even simple module layout parameter changes such as rotating selected modules, altering probe spacing, and staggering modules can improve published probe designs.

#### 3.3.1 Effect of optode orientation on probe characteristics

Re-orienting modules within existing probes alters the SDS distribution and, consequently, the probe’s *S_brain_*. In Fig. 7, we replicate the 36 mm^2^ *μ*NTS fNIRS module and its probe configuration described by Chitnis *et al.*^29^ The modules in the initial tessellation are oriented in the same direction as in the original paper (Fig. 7a). The spacing between each module is set to 5 mm and the *SDS_max_* is set to 30 mm to replicate the probe’s 4-module configuration. Each *μ*NTS module has 2 sources and 4 detectors, resulting in 8 intra-module channels per module ranging from 8 to 29 mm. The intra- and inter-module channel distribution and channel count resulting from the MOCA analysis of this particular probe are shown in Fig. 7b.

**Fig 7.**
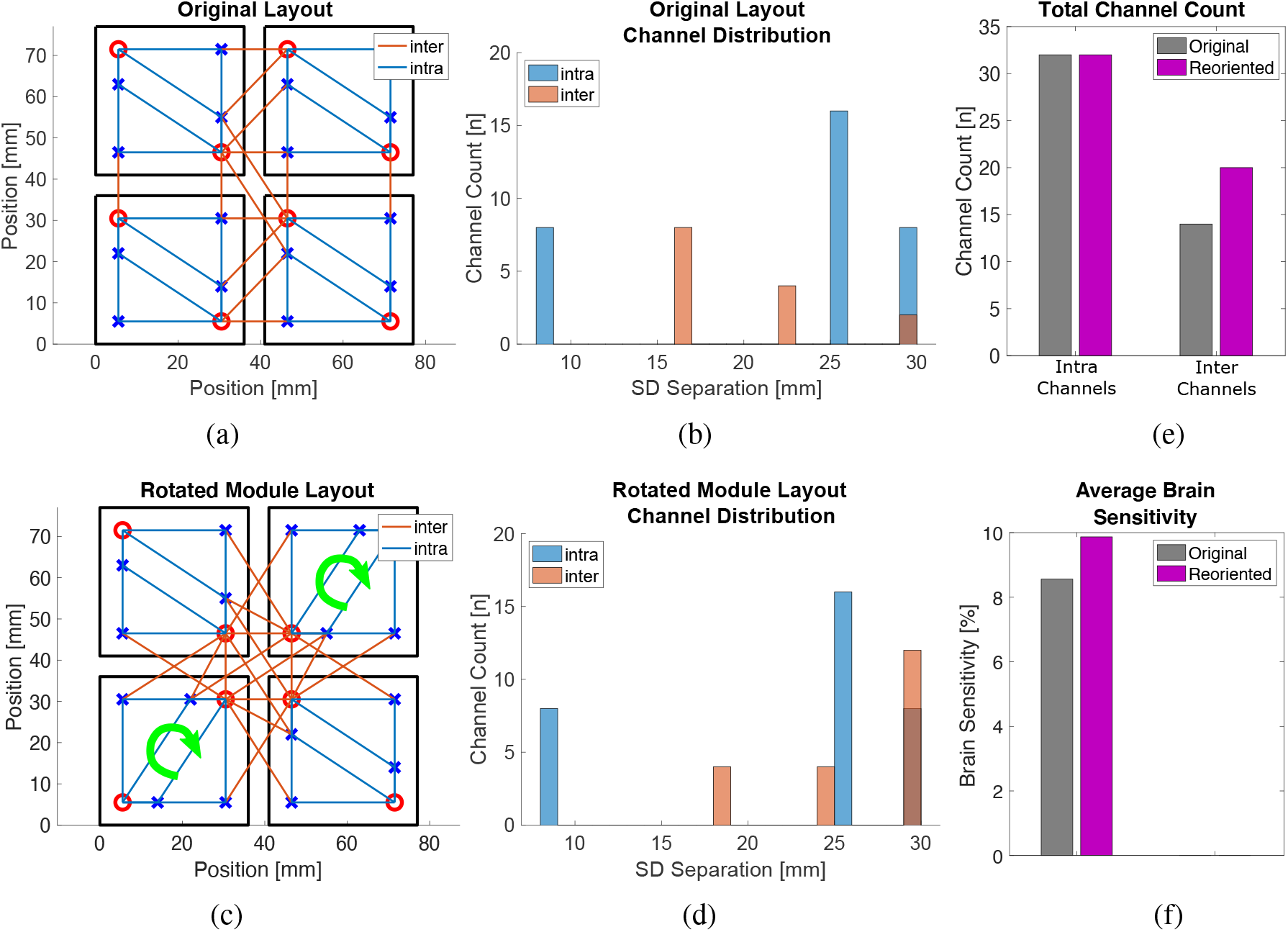
The 4-module probe based on the 36 mm2 *μ*NTS fNIRS module described by Chitnis *et al.*^29^ replicated using MOCA. (a) All modules are oriented in the same direction as the published probe, resulting in the intra- and inter-module channel distribution seen in (b). Red circles represent sources and blue crosses represent detectors. (c) Bottom-left and top-right modules are rotated 90 degrees clockwise with respect to orientation in (a). (d) Channel counts resulting from the probe configuration in (c). In channel distribution histograms (b, d), intra- and inter-module channels are shown in blue and orange, respectively. Dark orange indicates overlapping histogram counts. (e) Bar graph showing the total channel counts broken down by intra- and inter-module channel. (f) The resulting average brain sensitivity for the original and re-oriented probe layouts.

Fig. 7c shows the same 4-module probe but constructed with the bottom-left and top-right modules rotated 90 degrees clockwise. Using MOCA, the spatial channel plot overlaid onto this re-oriented probe shows a denser coverage of the center of the ROI compared to the original probe layout. The channel count distribution of this re-oriented probe is shown in Fig. 7d. As expected, the intra-module channels in Fig. 7b and Fig. 7d are identical. However, re-orienting the two modules produces a shift towards longer separation inter-module channels that are known to be more sensitive to brain tissues. The number of inter-module channels within the 10 to 20 mm range decreases from 8 to 4 and the number of 29 mm separation inter-module channels increases from 2 to 12 upon re-orienting the 2 modules. The re-orientation of modules not only allows the probe to have more long-separation channels, it also increases the total number of inter-module channels from 14 to 20 (Fig. 7e). Additionally, 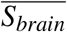 of the probe increases from 8.56% to 9.87% (Fig. 7f) while the number of spatial multiplexing groups, and subsequently the probe’s sampling rate, remains the same.

#### 3.3.2 Effect of probe spacing on probe characteristics

Probe spacing—the distance between edges of adjacent modules in a probe—is a parameter that can vary the resulting channel distribution and channel density of a probe by altering the relative distances between optodes on neighboring modules. To investigate the effect of this parameter, in Fig. 8, we replicate the probe layout described by Zhao *et al.*,^33^ which utilizes hexagonal shaped LUMO fNIRS modules developed by Gowerlabs.^63^ The length of each side of a LUMO module is set to 18 mm and each module contains three sources and four detectors. The *SDS_max_* is set to 30 mm. A uniform spacing is set between all adjacent modules. Here, we vary probe spacing from 0 to 15 mm in 5 mm increments.

**Fig 8.**
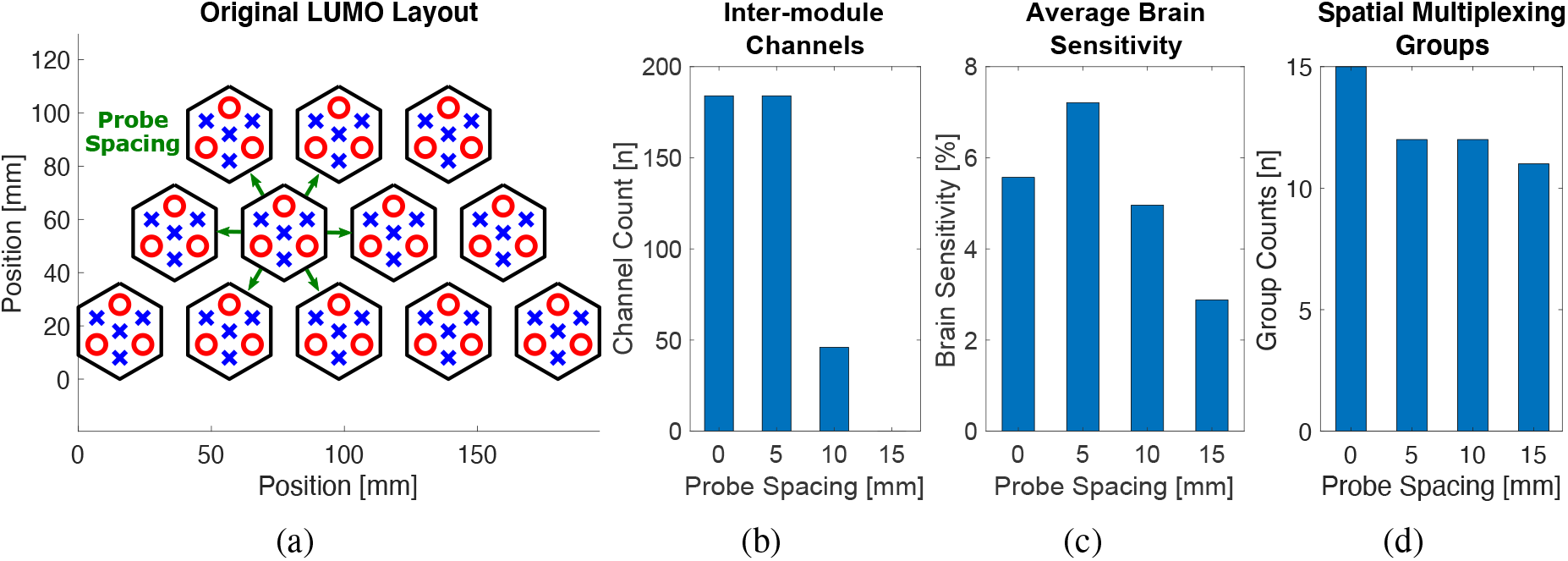
The hexagonal LUMO modules in a twelve-module probe described by Zhao *et al.*^33^ (a) Green arrows indicate the distances between modules that vary as probe spacing varies. (b), (c), and (d) show the inter-module channel count, average brain sensitivity, and the number of resulting spatial multiplexing groups, respectively, at probe spacing values of 0, 5, 10, and 15 mm. LUMO module orientations are held constant.

When all modules are densely packed with a spacing of 0 mm, the probe results in 184 intermodule channels, an 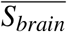 of 5.57%, and 15 SMGs. When the probe spacing is increased to 5 mm, the number of inter-module channels remains the same (Fig. 8b), 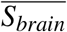 increases to 7.21% (Fig. 8c), and the *n_SMG_* decreases to 12 (Fig. 8d). The increase in 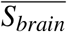 arises due to the overall increased distances between sources and detectors of inter-module channels which sample deeper into the brain tissue. At the same time, the increase in probe spacing increases the relative distance between adjacent sources, allowing more sources to be turned on at the same time and decreasing the *n_SMG_* needed.

When we increase probe spacing to 10 mm, the inter-module channel separations increase to above the *SDS_max_*. This decreases the number of “usable” inter-module channels and the probe’s 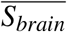. The *n_SMG_* remains unchanged between 5 and 10 mm probe spacing. This trend continues as we increase probe spacing to 15 mm at which point the number of inter-module channels drops to zero (Fig. 8b). Consequently, the probe’s 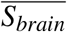 reaches a minimal plateau of 2.88% at 15 mm spacing because only intra-module channels above the SS threshold remain within the SD range (Fig. 8c). Similarly, since modules are farther apart, the *n_SMG_* drops to 11 (Fig. 8d).

#### 3.3.3 Effect of staggering modules on probe characteristics

Staggering of adjacent modules (Fig. 9f) within a high-density probe allows inter-module channel separations to increase due to the increased separation between neighboring optodes. To demonstrate the effect of staggering on the resulting probe, we replicate the M3BA module and its tessellation described by Von Lühmann *et al.*^30^ in Fig. 9. Each 42 mm^2^ M3BA module contains two sources and two detectors.

**Fig 9.**
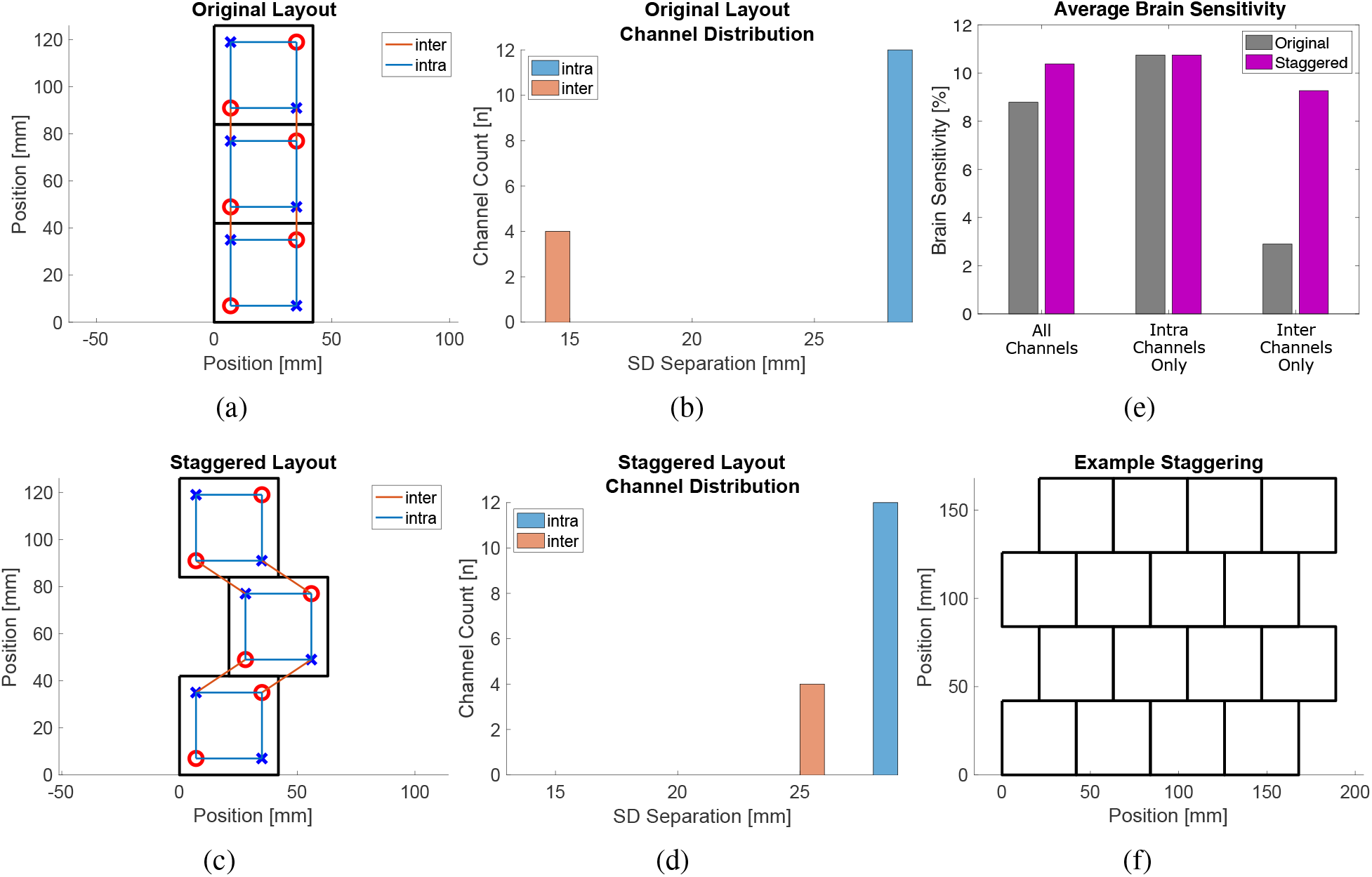
The M3BA modules described by Von Luühmann *et al.*^30^ replicated in MOCA. (a) M3BA modules in the published three-module tessellation. Red circles represent sources and blue crosses represent detectors. (b) The resulting intra- and inter-module channel distribution from probe layout in (a). (c) The staggered layout using the same three M3BA modules and (d) the resulting channel distribution. (e) Comparison of the average brain sensitivity for each of the two probe layouts, (a) and (c), separated by intra- and inter-module channel contributions. (f) An example 16-module probe with the second and fourth rows from the bottom staggered.

In Fig. 9a, we overlaid the intra- (blue) and inter-module (orange) channels over the three-module probe. The resulting channel distribution shows 12 intra-module channels at 28 mm and 4 inter-module channels at 14 mm SD separations (Fig. 9b). The 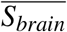 of this probe using all channels is 8.79% (Fig. 9e). When analyzed separately by intra- and inter-module channels, the 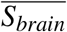 using only intra-module channels (10.75%) is larger the 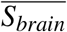 when using only inter-module channels (2.9%) since in this tessellation intra-module channels are larger and probe deeper into the tissue (Fig. 9e).

In Fig. 9c, we stagger the tessellated module layout by translating the center module by half of the module’s dimension in the horizontal axis. This alteration increases the inter-module channel separations to 25 mm (Fig. 9d). Consequently, the 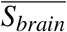 due to only inter-module channels increases to 9.27% (Fig. 9e). The 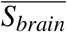 using all channels increases from 8.79% in the original tessellation to 10.38% in the staggered tessellation. The *n_SMG_* between the two layouts remained the same at 4.

## 4 Discussion

The design and analysis of modular fNIRS probes can quickly become complex due to the large number of module- and probe-scale parameters that must be taken into account. Here, we use MOCA to analyze full-head probes composed from vastly different elementary module shapes with the same optode layout. We also investigate the effect of module re-orientation, spacing between modules, and module staggering to improve existing fNIRS probes.

Fig. 4 reveals that, despite having the same optode layout, probes composed of different module shapes covering the same ROI result in different SD separation distributions. Although the intermodule channels are identical between modules, the resulting total number of channels is related to the number of modules needed to cover the ROI. The effect of module shape on channel distribution is complex and requires a tool like MOCA to thoroughly investigate. Certain module geometries result in optodes closer to the module’s edges, effectively shortening inter-module channels of completely tessellated probes. Because the optode layout in Fig. 3 is not completely symmetric and each module shape is an equilateral polygon, each individual module can be re-oriented without overlapping while maintaining the complete tessellation of the probe. While not altering intra-module channel distributions, these orientation configurations spatially alter channel locations and alter inter-module channel separations. The results from Fig. 4 also show how some individual module shapes may be more appropriate for subject populations. For example, the high count of 19 mm inter-module channel separations of the hexagon-based probe make it better suited for infant populations. These results demonstrate the dependency a probe’s derived characteristics have on module shape even when different modules have the exact same optode layout.

The results in Fig. 5 provide a counter-example where more channels or higher channel density may not necessarily lead to a more “effective” probe design. Despite having fewer total channels than the hexagon-based probe, the square-based probe results in a higher average brain sensitivity 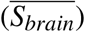 due to larger inter-module channel separations. This emphasizes the need for *S_brain_* to be considered in conjunction with channel distribution when comparing probes. Additionally, this analysis reveals that the use of inter-module channels in addition to intra-module channels does not always lead to a more efficient probe for different module shapes. In fact, the use of only inter-module channels increases the average penetration depth for the square- and triangle-based probes due to the larger channel separations. For the hexagon-based probe, however, Fig. 5 demonstrates that the contribution to 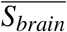 from using only intra- or only inter-module channels differed by merely 0.1%. Thus, users of the hexagon-based probe may benefit from the simplicity of using only intra-module channels rather than implementing a potentially complex data acquisition method to capture inter-module channels. These results show that it should not be assumed that adding inter-module channels to intra-module probes will always improve 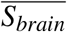.

Fig. 6 indicates that the hexagon-based probe can achieve the highest sampling rate among the three configurations if a spatial multiplexing encoding strategy is implemented. The frame rate of a sequential encoding strategy is dependent on the total number of sources (*n_s_*) because each source needs to be turned on and sampled once. Spatial multiplexing allows multiple sources within a group to be turned on simultaneously, allowing the sampling rate to increase by a factor of *n_s_/n_SMG_*, defined as the spatial multiplexing ratio (SMR). Therefore, despite having the lowest sampling rate when sampled sequentially due to the highest *n_s_* (Table 2, Row 1), the hexagon-based probe has the fastest sampling rate of the three probes when spatial multiplexing is used due to the low *n_SMG_* (Fig. 6a). These results demonstrate that a probe’s sampling rate can be increased by using different module shapes with the same optode layout.

Additionally, altering a probe’s tessellation can also improve a probe’s characteristics. Fig. 7 shows how existing probes can improve 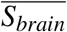 at no increased cost and without re-designing the module by optimizing the orientations of the modules. Such orientation optimization not only increases the channel density at the center of the ROI, but also increases the number of inter-module channels by 43%. The emerging inter-module channels also have larger source-detector separation (SDS) and contribute to an increase in 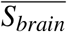. The re-oriented probe in Fig. 7 is only a representative case of how the *μ*NTS modular probe can be improved and is by no means exhaustive. It benefits from the asymmetry of the optode layout within each module. If the optode layout was symmetric, re-orienting modules would have no effect on either inter- or intra-module channels.

In Fig. 8, we investigated the effect of the spacing between modules on the derived characteristics of a probe based on LUMO modules.^33^ The results suggest that varying module spacing does have an impact on 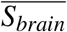. Since optodes are generally placed near the edges of the modules to maximize intra-module channel separations, dense probes with modules near one another tend to have shorter inter-module channel separations. This trend becomes more apparent as the size of the module increases. Increasing the probe spacing increases the distance between optodes on neighboring modules, thus increasing the 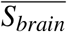 in the process. This increase in 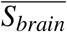, however, has a local maximum. As shown in Fig. 8, further increasing probe spacing leads to a drop in the number of inter-module channels as their SD separations become greater than the separation limit (*SDS_max_*). Additionally, increasing the distance between modules reduces the number of multiplexing groups (*n_SMG_*). Once the probe spacing exceeds the user-specified SMG diameter, one source on each module can be turned on at the same time because each source would be outside each other’s “effective” region. Because the distance between sources on the same module does not change, a different SMG is required for each source within a module. Thus, the limit to the minimum *n_SMG_* is equal to the number of sources on a single module. Fig. 8 shows that probe spacing can both alter *n_SMG_* to help meet sampling rate requirements and alter inter-module channel separations to meet channel distribution needs.

Fig. 9 shows that a staggering module layout can be used to increase 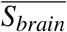 in dense probes. Dense coverage may prevent these probes from increasing probe spacing while their spatial restrictions limit potential module re-orientations. Simulations using a published module shape^30^ with zero probe spacing results in inter-module channels of 14 mm separations. These channels are too long to be short-separation (SS) channels and too short to be long-separation (LS) channels. Staggering such modules spatially increases inter-module channel separations while maintaining the compactness of a probe. For module designs with symmetrical optode layouts, we recommend staggering layouts by translating every other module row by half of the module’s maximum width in one axis (Fig. 9f). This ensures the optodes from the translated module are well separated from modules of the adjacent rows.

The results above are derived from investigating the module-scale and probe-scale design parameters that MOCA currently supports. However, this only represents a small subset of the general parameters previously used in evaluating a modular probe.^24^ For example, user feedback-based design parameters not yet accounted for in MOCA include conformability (a module’s ability to conform to a curved surface), subject comfort, and safety limits such as operating voltage and heating effects. Source output power and module weight each require external instrumentation measurements while noise-equivalent power and dynamic range calculations require lab-specific phantoms. Power consumption and a probe’s sampling rate depend on the type of optodes used as well as the control electronics of the individual module while a probe’s battery life can be adjusted using existing off-the-shelf components. Each of these design parameters are based on specific electronic or material components chosen for a particular module design. MOCA was built to easily scale and incorporate more complex mechanical-, ergonomic-, safety- and experiment-specific considerations in the future as those design parameters are evaluated.

There are limitations to MOCA’s current minimal subset of design parameters. First, the ability to re-orient, increase spacing, or stagger modules assumes that modules can be connected in any orientation. This is true for many published modular designs where cables of different lengths can be easily connected to the top of a module, but does not necessarily apply to more sophisticated designs that have embedded printed flex connectors that define inputs and outputs for modules. Second, MOCA does not currently support multiple module shapes within the same probe or different optode layouts on different modules. Third, MOCA’s channel count output does not include wavelength number as a multiplier. This approach allows one to quickly scale the channel distribution/counts when dual-wavelength or triple-wavelength sources are utilized. Similarly, the *n_SMG_* are also defined for a single wavelength, thus, when estimating the total sampling rate of the probe using multi-wavelength sources, the control unit must cycle through each group multiple times(once for each wavelength). Fourth, MOCA’s analysis is based on the coordinates of the center of an optode’s active area and does not account for the actual size of optode packages, the shape of the optode’s active area, or any master control unit needed to control a series of modules. Despite being able to place optodes near the edge of modules in MOCA, in practice, designers may face constraints imposed by the fabrication process due to board materials, sizes, and electrical routing needed to drive these optoelectronics. In general, shapes with large interior angles allow optodes to be placed closer to the module’s perimeter. Finally, MOCA currently outputs 2-D (flat) probe layout, and relies on other existing software, such as AtlasViewer,^41^ to perform 3-D head contour registration. In addition, the performance metrics outputted by MOCA are currently based on a 2-D probe layout and do not account for changes in SDS if the probe is “wrapped” on the 3-D surface of a head.^34^ Consequently, 2-D derived metrics may underestimate the number of channels when a probe is made to conform to the scalp. This may result in an increase in the number of total inter-module channels for a probe. We plan to address these limitations in the future improvement of MOCA.

## 5 Conclusion

We have developed a MATLAB-based toolbox – MOCA – with the goal of providing the fNIRS community with a systematic yet easy-to-use software platform to enable explorations of the large design space of module-based fNIRS probes. MOCA relies on both module-level parameters such as size, shape, and optode layout as well as probe-level parameters such as the maximum source-to-detector separation and ROI geometry to characterize a modular probe. It offers several convenient and automated functions, such as tessellating user-defined ROI and module connection routing optimization, to facilitate the design of a large modular fNIRS probe. It also outputs a set of informative fNIRS-relevant metrics, including channel distribution, average brain sensitivity, and source multiplexing group number, to allow quantitative characterization and comparison between various design decisions. In this work, we demonstrated, using representative examples, that a designer can utilize MOCA to optimize probes made of published modules without needing to re-design or re-fabricate existing modules, compare complex full-head probes composed of vastly different module shapes, and be used as a platform to systematically design future fNIRS modular designs.

## Disclosures

The authors declare no conflicts of interest.

## Acknowledgments

This research is supported by National Institutes of Health (NIH) under National Institute of Neurological Disorders and Stroke (NINDS) grant R01-EB026998 and National Institute of General Medical Sciences (NIGMS) under grant R01-GM114365.

## Code, Data, and Materials Availability

Our Modular Optode Configuration Analyzer (MOCA) toolbox is freely available for download at http://github.com/COTILab/MOCA.

**Morris Vanegas** is a doctoral candidate in Biomedical Engineering at Northeastern University. He received a BS in Aeronautical Engineering with Information Technology and dual MS in Aerospace and Mechanical Engineering from the Massachusetts Institute of Technology. His research interests are in portable and wearable near-infrared imaging systems.

**Dr. Miguel Mireles** is a postdoctoral research associate in the Biomedical Engineering Department at Northeastern University, Boston, USA. He received his PhD in Photonics from ICFO - The Institute of Photonic Sciences, Barcelona, Spain. His research interests span from the design and development of near-infrared systems to their application in neuroimaging and oncology treatment monitoring.

**Dr. Qianqian Fang** is currently an Associate Professor in the Bioengineering Department, North-eastern University, Boston, USA. He received his PhD from Dartmouth College in 2005. He then joined Massachusetts General Hospital and became an assistant professor in 2012, before he joined Northeastern University in 2015. His research interests include translational medical imaging systems, low-cost point-of-care devices for resource-limited regions, and high performance computing tools to facilitate the development of next-generation imaging platforms.

## Notes

### Competing Interest Statement

The authors have declared no competing interest.

